# Patterns of acquired HIV-1 drug resistance mutations and predictors of virological failure in Moshi, northern Tanzania

**DOI:** 10.1101/2020.04.21.052902

**Authors:** Shabani Ramadhani Mziray, Happiness H Kumburu, Hellen B Assey, Tolbert B Sonda, Michael J Mahande, Sia E Msuya, Ireen E Kiwelu

## Abstract

Drug resistance is a public health concern. Profiles of HIV drug resistance mutations (HIVDRM) and virological failure (VF) are not extensively studied in Tanzania. This study aimed to determine HIVDRM and predictors of VF in HIV-infected individuals failing first-line HIV drugs in Moshi, northern Tanzania. A case-control study was conducted at KCMC, Mawenzi, Pasua and Majengo health facilities with HIV-care and treatment clinics in Moshi from October, 2017 to August, 2018. Cases and controls were HIV-infected individuals with VF and viral suppression (VS) respectively. HIV-1 reverse transcriptase and protease genes were amplified and sequenced. Stanford University’s HIV drug resistance database and REGA HIV-1 Subtyping tool 3.0 determined HIVDRM and HIV-1 subtypes respectively. Odds ratios with 95% confidence intervals investigated predictors of VF. P-value <5% was considered statistically significant. A total of 124 participants were recruited, of whom 63 (50.8%) had VF and 61 (49.2%) had VS. Majority (66.1%) were females. Median [IQR] age and duration on ART were 45 [35-52] years and 72 [48-104] months respectively. Twenty five out of 26 selected HIV-1 RNA samples from cases were successively sequenced. Twenty four samples (96%) had at least one major mutation conferring resistance to HIV drugs, with non-nucleoside reverse transcriptase inhibitor (NNRTI) associated mutations as the majority (92%). Frequent NNRTI resistance mutations were K103N (n=11), V106M (n=5) and G190A (n=5). Prevalent nucleoside reverse transcriptase inhibitors mutations were M184V (n=17), K70R (n=7) and D67N (n=6). Dual-class resistance was observed in 16 (64%) samples. Thirteen samples (52%) had at least one thymidine analogue-associated resistance mutation (TAM). Three samples (12%) had T69D mutation with at least 1 TAM. Age was independently associated with VF [aOR 0.94 (0.90-0.97) p<0.001]. In conclusion, HIV drug resistance is common among people failing antiretroviral therapy and resistance testing will help to guide switching of HIV drugs.

## Introduction

Human Immunodeficiency Virus (HIV) infection is still a public health problem with an estimation of 37.9 million people living with it worldwide by the end of 2018. Among the people living with HIV (PLHIV) globally, 36.2 million are adults, 1.7 million are children below 15 years and 18.8 million are women aged 15 years and above. Every day, 5000 people are infected with HIV globally, and more than a half of these new infections occur in sub-Saharan Africa (sSA). Eastern and Southern Africa alone harbours more than a half of the world burden of HIV-infections with 20.6 million of PLHIV by the year 2018 [1]. The incidence-prevalence ratio (IPR) of the worldwide HIV-infection has declined considerably from 11.2% in 2000 to 4.6% in 2018. In the same year, sSA was also observing a decline in HIV-infection IPR to about 3.9% and 5.5% in Eastern and Southern Africa; and Western and Central Africa respectively. Despite the decline in IPR, HIV-infection is still a public health problem in sSA due to unacceptably high number of PLHIV in this setting (approximately 68% of the global burden) with substantial new HIV-infections [1].

Introduction of highly active antiretroviral therapy (HAART) in sSA countries including Tanzania has significantly helped to reduce HIV/AIDS related mortality from around 900,000 in 2010 to 470,000 in 2018 [1] with improved quality of life [2]. The globe further committed that by the year 2020, 90% of PLHIV should know their HIV status, 90% of PLHIV who know their HIV status should be on ART, and also 90% of PLHIV on ART should enjoy the sustainable viral suppression, this is alias known as 90-90-90 target [3]. In the race to achieve the second UNAIDS ‘90’ target, about 51% of PLHIV are on HAART in Western and Central Africa (WCA); and 67% in Eastern and Southern Africa (ESA) respectively. The proportions of HIV-infected individuals on treatment with an enjoyment of viral suppression is better in WCA (79%) compared to ESA (58%) respectively, however, the numbers are below the third UNAIDS ‘90’ target [1]. One of the key challenges which partly explain the failure to achieve the third UNAIDS ‘90’ in sSA is the emergence of HIV drug resistance (HIVDR).

HIVDR is defined by World Health Organisation (WHO) as presence of one or more mutations in HIV drug targeted genes that compromise the ability of a specific drug or combination of drugs to block replication of HIV [4]. HIVDR reverses the gains of HAART in HIV-infected individuals on treatment for at least 6 months. HIV-infected individuals on failed first-line HAART regimens are reported to have 50 to 97% of non-nucleoside reverse transcriptase inhibitors (NNRTI) resistance world-wide [4]. In sSA, more than 80% of HIV-infected individuals with VF on first-line HAART have HIV-drug resistance [5]. In addition, sSA has been reported to accommodate tenofovir resistance in more than a half of people with first-line HAART failure on tenofovir-based regimens. Cytosine analogue resistance was also highly evident in sSA as compared to Western Europe. Eastern Africa was more commonly reported to have lamivudine and emtricitabine resistance than NNRTI resistance [6].

Tanzania introduced antiretroviral therapy (ART) program in 2004. As of 2016, the total number of people on ART was 839,544, and the current recommended first-line HAART for adults and adolescents is tenofovir + lamivudine + dolutegravir. In case of first-line HAART failure, the recommended second-line HAART for adults and adolescents is zidovudine/lamivudine + ritonavir-boosted atazanavir or tenofovir/emtricitabine + ritonavir-boosted atazanavir [7]. In Tanzania, the markers for treatment failure are based on immunological, clinical and WHO recommended virological criteria [8]. Clinical and immunological criteria were extensively described to be less sensitive and less effective [9] and may reduce early notifications of VF. Tanzania continues to scale up HIV viral load (HVL) testing which is currently done at few selected settings. Of more public health importance is that HIV-infected individuals confirmed to have VF are switched to second-line HAART without programmatic HIV drug resistance testing and monitoring, a practice which may transfer cross-resistance patterns to newly switched drugs and limit the choices for few available treatment options.

Recently, few studies in Tanzania have reported a wide range of first-line HIV drug resistance mutations (HIVDRMs) including the thymidine analogue associated mutations which compromise susceptibility to multi-NRTIs [10–12]. In order to complement the efforts to unearth the burden of HIVDR and preserve the integrity of limited second-line HIV drugs in Tanzania, this study aimed to test genotypically for drug resistance in the reverse transcriptase (RT) and protease genes from individual with VF on first-line HAART in Moshi municipality, northern Tanzania. The study also explored the independent predictions for VF.

## Materials and methods

### Study design and settings

Between October, 2017 and August, 2018, unmatched case-control study was done in four HIV/AIDS care and treatment clinics (CTC) in Moshi municipality, situated in Kilimanjaro region, northern Tanzania. Moshi municipality is one of the seven districts of Kilimanjaro region. The municipality has a total of 18 health facilities with CTC. Four out of 18 CTCs with high client volume were purposively selected to participate in the study. The CTCs included were that of Kilimanjaro Christian Medical Centre (KCMC), Mawenzi regional referral hospital, Majengo and Pasua health centres. KCMC provides tertiary care hospital services to around 6.8 million people living in the northern zone of Tanzania (Tanga, Kilimanjaro, Arusha and Manyara) and other referrals from nearby settings. Mawenzi regional referral hospital provides referral medical services to around 1.6 million people in Kilimanjaro region.

At the time of recruitment of study participants, the management of HIV-infected individuals was done in-line with national guidelines for management of HIV and AIDS of 2017 [8]. The CTCs routinely provided HIV counselling and testing services, ART care and treatment, treatment monitoring, laboratory investigations and treatment adherence support to HIV-infected individuals. The recommended first-line HAART for adults and adolescents at the time of the study was tenofovir + lamivudine + efavirenz (TLE), given as a single dose formulation. Prescriptions of antiretroviral drugs were done by clinicians and to some points by trained nurses in the health centres. HVL testing was routinely done to all HIV-infected individuals who have been on ART for at least 6 months. Those found with HVL > 1,000 copies/ml were offered enhanced adherence counselling (EAC) by trained health care providers. EAC was provided on monthly bases for three months at the CTCs. After 3 months of EAC, retesting of HVL was done and HIV-infected individuals found still to have HVL > 1,000 copies/ml were classified as having VF and followed by further actions to change their regimens to second-line. The clinic visits for adolescents and youths were scheduled at different times from adults to maximize adherence counselling and support. Adherence level of ≥ 95% and < 95% was regarded as good and poor respectively [8].

### Study population

The study population was HIV-infected individuals attending the selected CTCs for routine care and were on first-line HAART treatment for a year or more. HIV-infected individuals with VF and viral suppression (VS) were included into study as cases and controls respectively. The cases were defined as HIV-infected individuals with >1000 copies/ml of HIV plasma RNA confirmed by <0.5 logarithmic difference between initial and second HVL with EAC in between. Controls were the HIV-infected individuals attending the same CTCs along with cases but they have viral suppression (HVL < 1000 copies/ml). The cases found to have less than three EAC were excluded from the study. The study further excluded the cases with significant difference HVL measured at the date of interview and that measured before starting EAC (HIV-1 plasma RNA viral log_10_ drop greater than 0.5 at three month interval with 3 EAC in between).

### Sample size determination

We assumed that the study had 80% power and the proportion exposed in the control group was 20%. With equal number of cases and controls (ratio of 1:1), we could manage to detect odds ratio (OR) of 3.0 or greater with the following sample size calculation as described by Charan and Biswas [13].

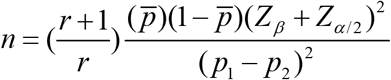

*n* = sample size in each group, *r* = ratio of controls to cases, 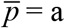 measure of variability (average proportion exposed), *p*_1_ = proportion exposed in cases, *p*_2_ = proportion exposed in controls, *p*_1_ − *p*_2_ = Effect size (the difference in proportions), *Z*_*α*/2_ = level of two-tailed statistical significance, *Z_β_*= standard normal variate for power of the study. Our study power was 80%, *Z_β_*= 0.84; Level of significance was 0.05, *Z_α_* = 1.96; *r* = 1 (equal number of cases and controls), *OR* = 3.0 and *p*_2_ = 20% or 0.2.

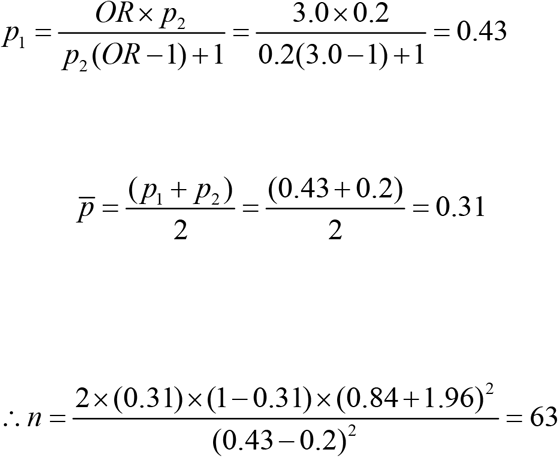

Therefore, the study planned to recruit 63 study participants with first-line VF and 63 with VS making a total sample size of 126.

### Enrolment and study procedures

Study participants who met the inclusion criteria and consented to participate were interviewed to collect demographic and clinical information. After the interviews, 8-10 ml of EDTA whole blood was collected and centrifuged at 2200 revolutions per minute for 10 minutes with brake-offs to separate plasma from buffy coat and red blood cells within 4 hours post-collection. HVL was enumerated in-vitro from plasma by reverse transcription-polymerase chain reaction as per Abbott m2000rt system and was expressed in copies/ml of plasma.

Due to financial constraints, 26 out of 63 samples of cases were selected for sequencing. Selection was based on having HVL > 14,000 copies/ml. Plasma HIV-1 RNA was extracted using PureLink^®^ Viral RNA/DNA Mini Kit (Invitrogen, Thermo Fisher Scientific, USA) as per manufacturer’s instructions. RT and protease genes of the extracted HIV-1 RNA was reversely transcribed into complementary DNA (cDNA) and subsequently subjected into nested polymerase chain reactions (PCR) according to manufacturer’s instructions prescribed in HIV-1 genotyping kit: Amplification module (Applied Biosystems, Life Technologies, Warrington, UK). The amplified cDNA was purified using ExoSAP-IT^™^ PCR product clean-up reagent (Applied Biosystems, Thermo Fischer Scientific, Inc.). Reactions were performed using HIV-1 genotyping kit: Cycle sequencing module (Applied Biosystems, Life Technologies, Warrington, UK) based on Sanger sequencing method using BigDye^™^ Terminator v3.1 cycle sequencing kit (Applied Biosystems, Thermo Fischer Scientific, Inc.). Sequencing was done using 3500xl genetic analyzer (Applied Biosystems) with a 24 capillary 50 cm array. The study profile was described in Fig **1**.

**Fig 1.**
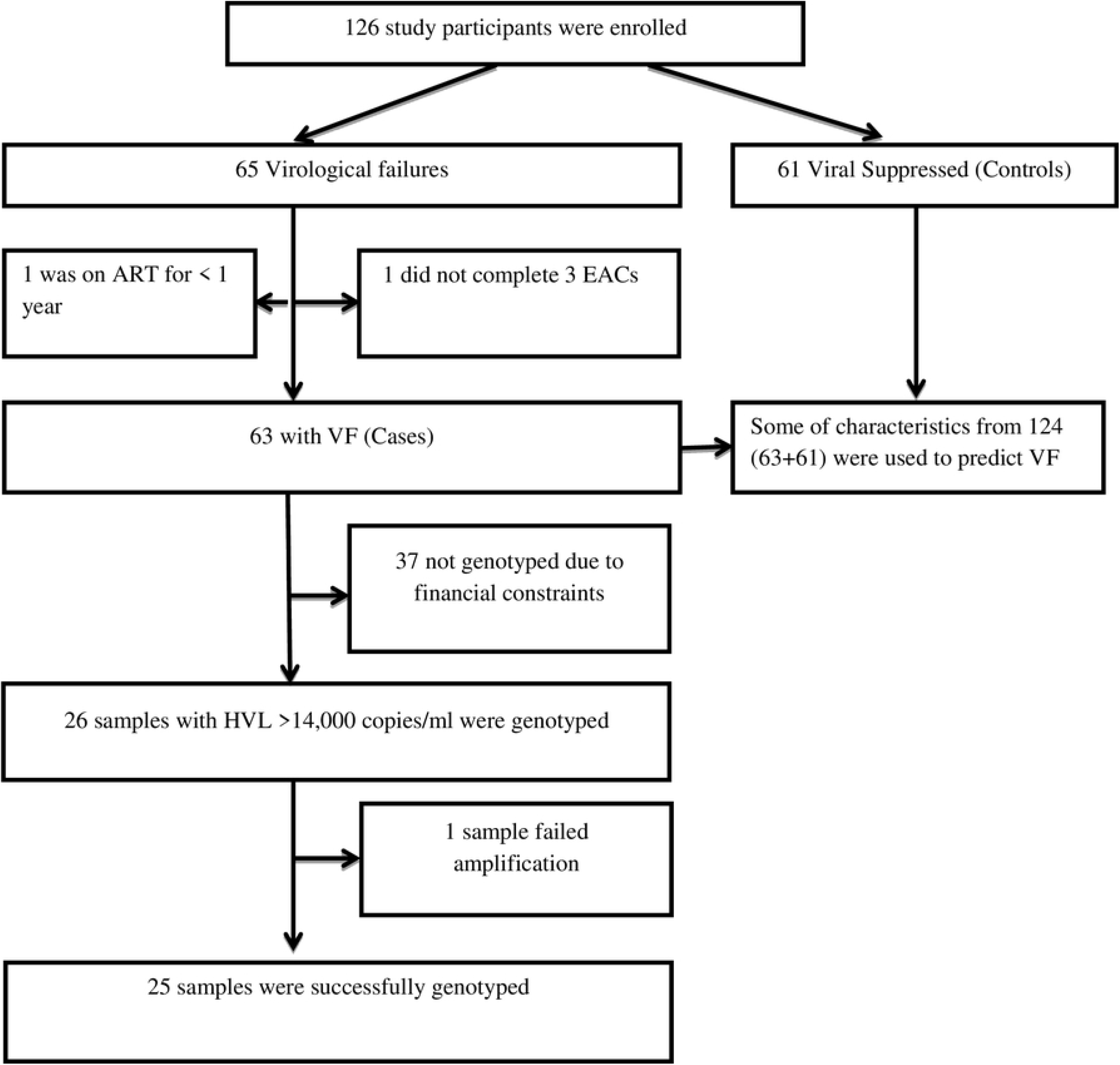
Study profile

### Data analysis

#### Sequence data analysis

The raw sequence data were assembled, aligned and edited using automated base calling software, RECall, available at (http://pssm.cfenet.ubc.ca) to generate individual consensus sequences. The sequence data were submitted to GenBank and obtained accession numbers MT347616 – MT347640. HIVDRMs were analysed using HIV drug resistance database of Stanford University available at (http://hivdb.stanford.edu/pages/algs/sierra_sequence.html) and HIVDRM mutation list from 2019 updates of drug resistance (AIDS Society). To get HIV-1 subtypes, generated consensus sequences were analysed using REGA HIV-1 Subtyping tool 3.0 available at (http://dbpartners.stanford.edu:8080/RegaSubtyping/stanford-hiv/typingtool/). REGA confirmed a subtype after clustering with a pure subtype in a database by > 800 base pairs with a bootstrap confidence of more than 70% in absence of recombination in the boot scan.

#### Statistical analysis

Data analysis was performed using STATA version 14.0 (College Station, Texas 77845-4512, USA). Categorical variables were summarized using frequency and proportion while mean or median with their respective measure of dispersion were used to summarize numerical variables. A Chi-square test was used to compare the differences in proportions between groups. Odds ratios (OR) and 95% confidence interval (CI) for predictors of virological failure were estimated using multivariable logistic regression model. A p-value <0.05 (2 tails) was considered statistically significant.

### Ethics Consideration

The study was approved by the Kilimanjaro Christian Medical Research Ethics Review Committee in 2017 with ethical clearance certificate number 2028. Approvals to conduct the study were further granted by respective authorities in four study sites. Patient information sheet was given to every potential study participant to ensure informed consenting process before and after commencement of the study, potential participants discussed it with the researcher. The sheet explained in detail the aim of the study, purpose, confidentiality, benefits, unconditional withdrawal from the study and risks of taking part into the study. Every study participant consented/assented in writing to take part in the study. The HIV drug resistance testing results were shared with the respective CTCs for further management of the HIV-infected individuals.

## Results

### Demographic characteristics of study participants

A total of 124 study participants were recruited in this study. Out of the 124, 63 (50.8%) of study participants had virological failure and 61 (49.2%) had viral suppression. The age of the 124 participants ranged from 15 to 79 years, with median [IQR] age of 45 [35-52] years. Of the 124; 82 (66.1%) were females and 89 (71.8%) were employed in either formal or informal sector (Table **1**).

**Table 1.** Demographic characteristics of study participants (N=124)

| Characteristic | Overall<br>n (%) | VF<br>n (%) | VS<br>n (%) |
| --- | --- | --- | --- |
| Age (years) |  |  |  |
| 15-34 | 29 (23.4) | 26 (41.3) | 3 (4.9) |
| ≥35 | 95 (76.6) | 37 (58.7) | 58 (95.1) |
| Median age [IQR] <sup>a</sup> | 45 [35-52] | 41 [21-49] | 48 [43-54] |
| Sex |  |  |  |
| Male | 42 (33.9) | 26 (41.3) | 16 (26.2) |
| Female | 82 (66.1) | 37 (58.7) | 45 (73.8) |
| Education level |  |  |  |
| Primary or none | 84 (67.7) | 36 (57.2) | 48 (78.7) |
| Secondary and above | 40 (32.3) | 27 (42.8) | 13 (21.3) |
| Marital status |  |  |  |
| Married/cohabiting | 40 (32.3) | 18 (28.6) | 22 (36.1) |
| Single | 84 (67.7) | 45 (71.4) | 39 (63.9) |
| Occupation |  |  |  |
| Employed | 89 (71.8) | 36 (57.1) | 53 (86.9) |
| Unemployed | 35 (28.2) | 27 (42.9) | 8 (13.1) |
| CTC enrolled |  |  |  |
| KCMC | 53 (42.7) | 38 (60.3) | 15 (24.6) |
| Majengo HC | 19 (15.3) | 7 (11.1) | 12 (19.7) |
| Mawenzi RRH | 29 (23.4) | 14 (22.2) | 15 (24.6) |
| Pasua HC | 23 (18.6) | 4 (6.4) | 19 (31.1) |

VF = Virological failure, VS = Viral suppression; <sup>a</sup> = Expressed as median [Interquartile range]

### Clinical characteristics of study participants

The median [IQR] time on ART for the 124 participants was 72 [IQR 48 - 104] months, 92 (74.0%) had good adherence, 65 (52.4%) were in non-tenofovir based HAART and 66 (53.2%) were in WHO clinical stage III/IV (Table **2**). The median CD4 count (cells/μL) at follow up was higher in VS participants 518 [IOR: 326-741] than in VF group 334 [IQR: 134-549]. Likewise 100% of VS participants reported good adherence to ART compared to 49% of VF participants (Table **2**).

**Table 2.** Clinical characteristics of the participants (N = 124)

| Characteristic | Overall | VF | VS |
| --- | --- | --- | --- |
|  | n(%) | n(%) | n(%) |
| Duration of ART intake (months) |  |  |  |
| 12 – 36 | 20 (16.1) | 7 (11.1) | 13 (21.3) |
| >36 | 104 (83.9) | 56 (88.9) | 48 (78.7) |
| Median [IQR] | 72 [48-104] | 84 [60-120] | 72 [48-96] |
| Adherence status |  |  |  |
| Good | 92 (74.2) | 31 (49.2) | 61 (100.0) |
| Poor | 32 (25.8) | 32 (50.8) | 0 (0.0) |
| Ever switched ART |  |  |  |
| Yes | 54 (43.5) | 30 (47.6) | 24 (39.3) |
| No | 70 (56.5) | 33 (52.4) | 37 (60.7) |
| HAART initiated |  |  |  |
| TDF based | 39 (31.5) | 12 (19) | 27 (44.3) |
| None TDF based | 85 (68.5) | 51 (81) | 34 (55.7) |
| HAART on use |  |  |  |
| TDF based | 59 (47.6) | 22 (34.9) | 37 (60.7) |
| Non TDF based | 65 (52.4) | 41 (65.1) | 24 (39.3) |
| Baseline CD4 count (cells/μL) | | | |
| <100 | 37 (29.8) | 17 (27) | 20 (32.8) |
| ≥100 | 87 (70.2) | 46 (73) | 41 (67.2) |
| Median CD4 count [IQR] | 173 [72-276] | 179 [68-256] | 169 [73-282] |
| Follow up CD4 count (cells/μL) | | | |
| <100 | 13 (10.5) | 12 (19) | 20 (32.8) |
| ≥100 | 111 (89.5) | 51(81) | 41 (67.2) |
| Median [IQR] | 444[215-628] | 334 [134-549] | 518 [326-741] |
| WHO clinical stage |  |  |  |
| I/II | 58 (46.8) | 26 (41.3) | 32 (52.5) |
| III/IV | 66 (53.2) | 37 (58.7) | 29 (47.5) |
| Time from HIV diagnosis to ART initiation (months)<sup>a</sup> |  | 3.5 [1.4 – 17.9] | 2.6 [0.9 – 14.0] |

**TDF**= Tenofovir Disoproxil Fumarate, **VF**= Virological failure **VS**= Viral suppression; **a**= Expressed as median [Interquartile range]

### Profiles of acquired HIV drug resistance in RT and protease genes

Genotyping of RT and protease genes of HIV-1 was successful on 25 out of 26 selected samples from participants with VF. Of the 25, almost all (n=24) had at least one major mutation conferring resistance to HIV drug (Table 3). Out of 25, 23 (92%) samples had at least one non-nucleoside reverse transcriptase inhibitor (NNRTI) resistance associated mutations (Fig **2**). The most frequent NNRTI mutations were K103N (44%), V106M (20%), and G190A (20%) (Fig **2**), all of these three mutations confer high level resistance to efavirenz and nevirapine.

**Table 3.** Characteristics of successfully genotyped samples (N=25)

| SN | Sample ID | Age (years) | Duration on HAART (years) | HAART at date of interview | Viral Load at date of interview | NRTI mutations | NNRTI Mutations | PI Mutations | HIV-1 Subtype |
| --- | --- | --- | --- | --- | --- | --- | --- | --- | --- |
| 1 | K03 | 18 | 10 | AZT+3TC+EFV | 389,006 | M184V, D67T, K70R,T215AITV, K219E | Y188L, K238T | None | A1 |
| 2 | K11 | 55 | 6 | ABC+3TC+LPVr | 24,463 | D67N, K70R, M184V,K219E, T215F | None | L10F,K20T, M46I,I47V, N88G | C |
| 3 | K12 | 29 | 6 | AZT+3TC+EFV | 20,805 | M184V | K103N,V108I, K238T | None | CRF 10_CD |
| 4 | K15 | 49 | 12 | AZT+3TC+NVP | 81,123 | None | None | None | A1 |
| 5 | K16 | 44 | 3 | TDF+3TC+EFV | 23,354 | None | K103N | None | A1 |
| 6 | K17 | 15 | 7 | AZT+3TC+NVP | 25,879 | None | G190A | None | A1 |
| 7 | K19 | 17 | 6 | ABC+3TC+EFV | 296,646 | M41L, D67N, K70R, M184V, T215Y, K219Q, L74I,T69D | A98G,K103N, P225H,F227F, 238T | None | C |
| 8 | K21 | 20 | 13 | ZVD+3TC+EFV | 33,887 | M184V,T215F | K103N,Y188F, M230L | None | C |
| 9 | K25 | 19 | 10 | AZT+3TC+EFV | 37,794 | None | V106M, G190A | None | C |
| 10 | K29 | 34 | 9 | AZT+3TC+EFV | 14,431 | None | K103S, V106M | None | C |
| 11 | K32 | 41 | 4 | TDF+3TC+EFV | 161,220 | A62V, K65R, M184V,K219E | K101E,V106M, Y181C, G190A | None | C |
| 12 | K36 | 29 | 10 | AZT+3TC+EFV | 23,056 | M184V | L100I, K103N | None | A1 |
| 13 | K37 | 15 | 9 | AZT+3TC+NVP | 20,540 | D67N, K70R, M184V,T215F, K219Q | A98G, K101E, G190A | None | C |
| 14 | K38 | 46 | 13 | ABC+3TC+ATVr | 1,635,827 | M41L, E44D, D67N, T69D, M184V,L210W, T215Y, K219R | K101E, E138K, Y181C, G190A, H221Y | L24I, L33F, M46I, I54V, Q58E,V82A , N88S | CRF 10_A1D |
| 15 | M02 | 35 | 5 | AZT+3TC+NVP | 124,714 | D67N, T69D, K70R, M184V | G190A | None | D |
| 16 | M03 | 69 | 9 | AZT+3TC+NVP | 36,600 | M41L, E44A, M184V,L210W, T215Y, K219N | Y181C, H221Y | None | A1 |
| 17 | M05 | 53 | 10 | AZT+3TC+NVP | 65,638 | D67G, K70R, M184V,T215FV, K219E | K103S, E138Q | None | A1 |
| 18 | M07 | 52 | 4 | ABC+3TC+EFV | 66,486 | M184V | K103N | None | A1 |
| 19 | M10 | 41 | 8 | AZT+3TC+NVP | 45,819 | D67N, K70R,<br>M184V,K219Q | K103N, V179L,<br>K238T | None | A1 |
| 20 | MJ01 | 48 | 9 | AZT+3TC+EFV | 14,065 | M184V,L210W, T215Y | K103N | None | C |
| 21 | MJ03 | 36 | 9 | TDF+3TC+EFV | 176,378 | None | K103N | None | C |
| 22 | MJ05 | 45 | 5 | TDF+FTC+EFV | 222,227 | None | L100I, K103N | None | C |
| 23 | MJ06 | 51 | 4 | TDF+3TC+EFV | 14,711 | K70E, M184V | K103N,V108I,<br>H221Y, F227L | None | D |
| 24 | P02 | 54 | 8 | AZT+3TC+EFV | 235,443 | None | V106M,V179D | None | C |
| 25 | P03 | 45 | 5 | TDF+3TC+EFV | 128,781 | M41L, K70S, L74I, M184V,<br>T215Y | V106M,E138Q,<br>F227L | None | C |

**Fig 2.**
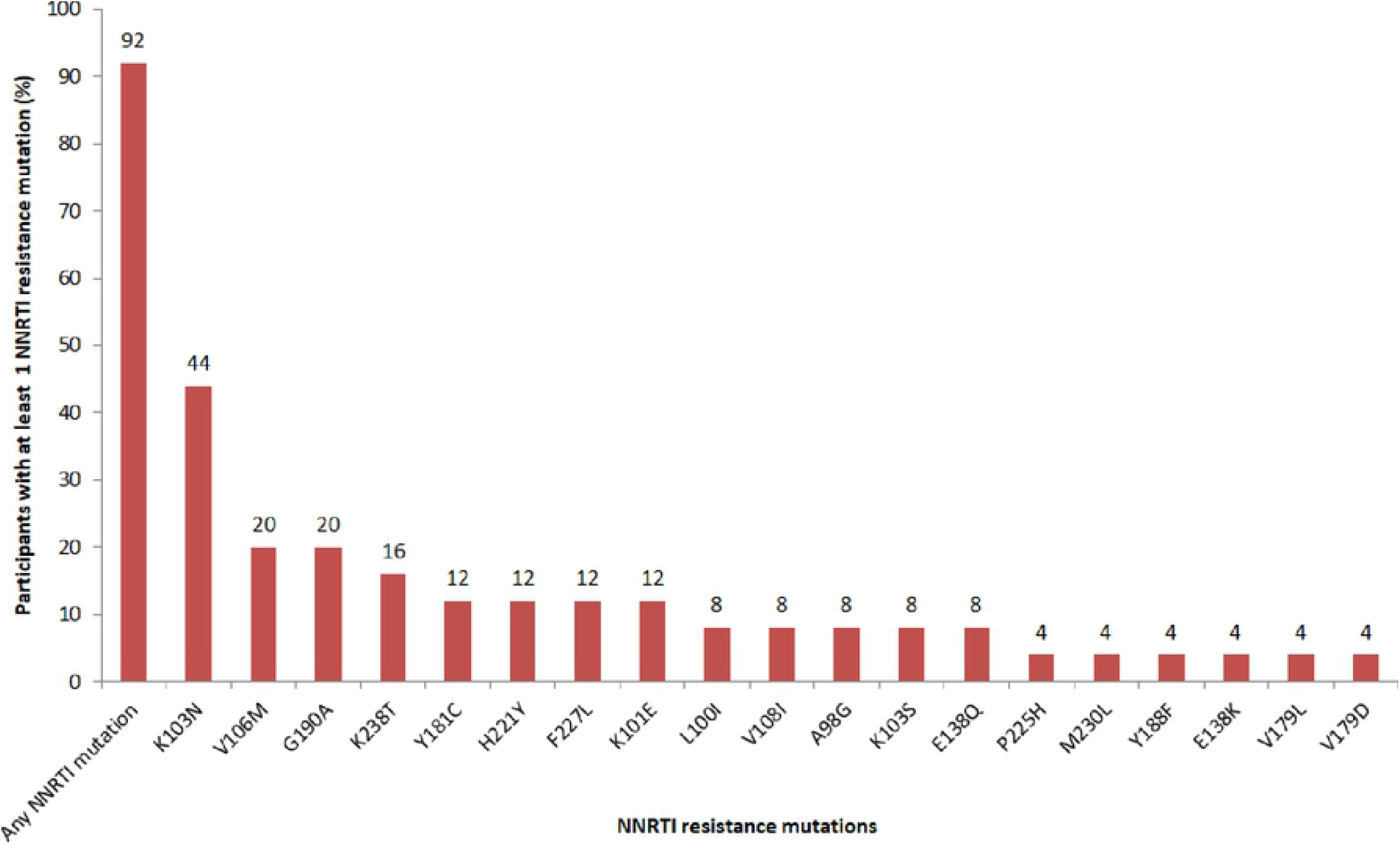
Profiles of NNRTI resistance mutations in participants with virological failure

In addition, 17 samples (68%) had at least one mutation associated with nucleoside reverse transcriptase inhibitors (NRTI) resistance. The most frequent NRTI mutations were M184V (68%), K70R (28%), and D67N (24%) (Fig **3**). M184V mutation was associated with high level resistance to emtricitabine, and lamivudine; while K70R and D67N are among thymidine analogue-associated resistance mutations (TAM) reducing the susceptibility of all current NRTI on use.

**Fig 3.**
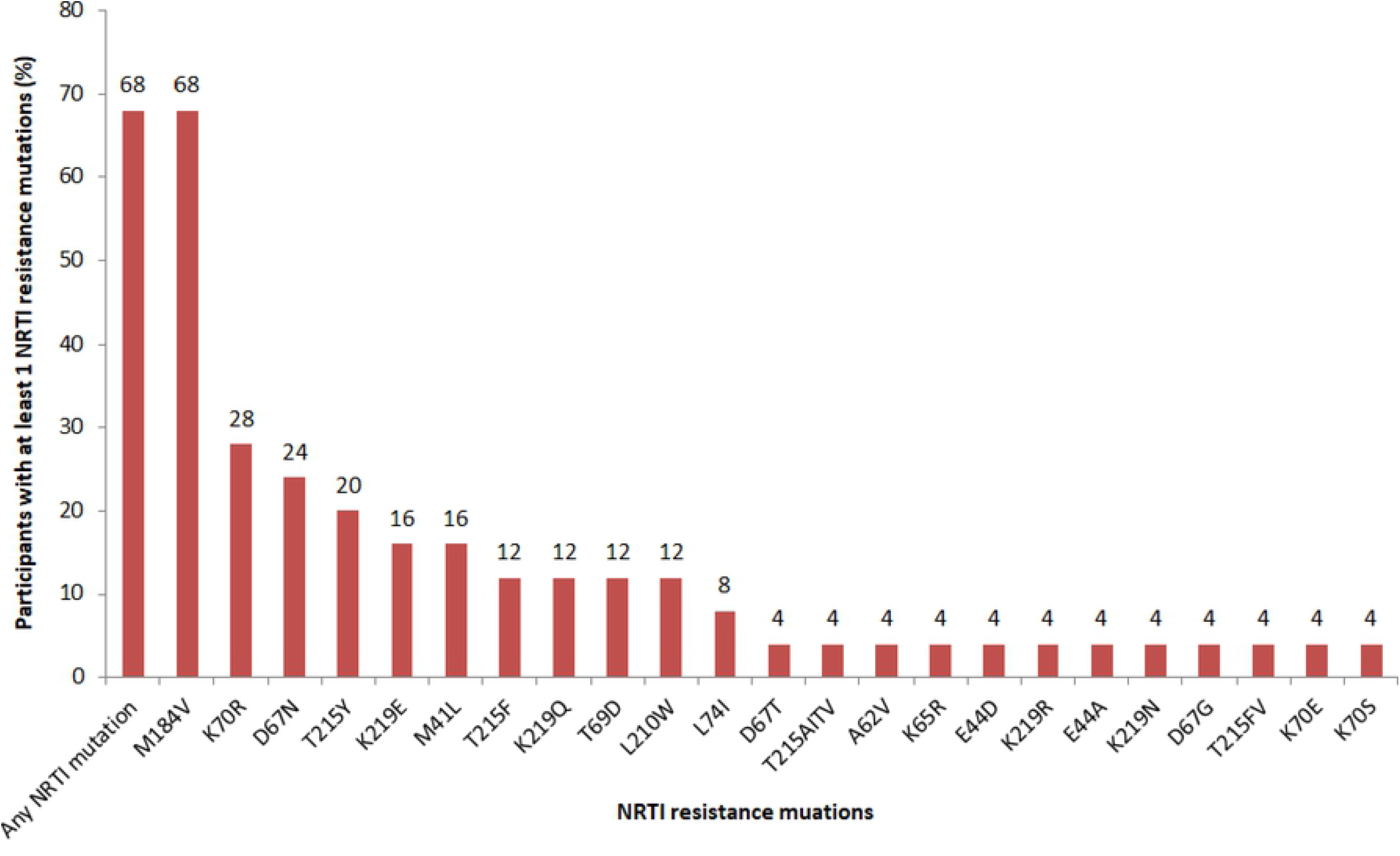
Profiles of NRTI resistance mutations in participants with virological failure

### Characteristics of successfully genotyped samples

The characteristics of successfully genotyped samples have been displayed in Table 3. Dual class resistance was observed in 16 (64%) samples. In general, thirteen samples (52%) had at least one TAM while three samples (12%) had T69D mutation together with at least 1 TAM. Identified TAMs were T215YF (n=9), K219QE (n=7), K70R (n=7), D67N (n=6), M41L (n=4), and L210W (n=3). Presence of T69D mutation along with at least 1 TAM reduces the susceptibility of all currently NRTIs on use. Of more scientific interest is that one sample was fully susceptible (no detectable HIV drug resistance mutation) despite having VF and substantial high HVL at the date of interview.

### HIV-1 diversity in the RT and protease gene

The HIV-1 diversity in the RT and protease genes is shown in Table **4**. HIV-1 subtype C was the most prevalent subtype (48%) followed by A1 (36%), D (8%) and recombinants (8%). The recombinants identified were A1D and CD. All the recombinants were circulating recombinant forms (CRF).

**Table 4.** HIV-1 diversity in RT and protease genes (N=25)

| Rega assignment | Number of sequences | Percentage |
| --- | --- | --- |
| HIV-1 Subtype C | 12 | 48 |
| HIV-1 Subtype A (A1) | 9 | 36 |
| HIV-1 Subtype D | 2 | 8 |
| Recombinant of A1, D | 1 | 4 |
| Recombinant of C, D | 1 | 4 |
| **Total** | **25** | **100** |

### Predictors of virological failure

The predictors of virological failure (VF) are shown in Table **5**. In bivariate analysis, participant age, occupation, HAART initiated and HAART on use were significantly associated with virological failure. In the multivariable logistic regression analysis, only age remained to be independent predictor of VF. We found that, one unit increase in participant age (year) was associated with 6% lower odds of VF [aOR 0.94 (0.90-0.97) p<0.001].

**Table 5.** Predictors of HIV virological failure (VF) (N=124)

| Characteristic | VF |  | Crude analysis |  | Multivariable analysis |  |
| --- | --- | --- | --- | --- | --- | --- |
|  | No (%) | Yes (%) | cOR (95% CI) | p-value | aOR (95% CI) | p-value |
| Age (years)* | 48 [43-54] | 41 [21-49] | 0.93 (0.90-0.96) | <0.001 | 0.94 (0.90-0.97) | 0.001 |
| Sex |  |  |  |  |  |  |
| Male | 16 (26.2) | 26 (41.3) | 1 |  | 1 |  |
| Female | 45 (73.8) | 37 (58.7) | 0.51 (0.24-1.08) | 0.079 | 0.56 (0.21-1.51) | 0.254 |
| Marital status |  |  |  |  |  |  |
| Married | 22 (36.1) | 18 (28.6) | 0.71 (0.33-1.51) | 0.373 | - | - |
| Single | 39(63.9) | 45 (71.4) | 1 |  |  |  |
| Occupation |  |  |  |  |  |  |
| Employed | 53 (86.9) | 36 (57.1) | 0.2 (0.08-0.49) | <0.001 | 0.35 (0.12-1.04) | 0.059 |
| Unemploye | 8 (13.1) | 27 (42.9) | 1 |  | 1 |  |
| d |  |  |  |  |  |  |
| Duration of ART intake (months) |  |  |  |  |  |  |
| 12 - 36 | 13 (21.3) | 7 (11.1) | 1 |  |  |  |
| >36 | 48 (78.7) | 56 (88.9) | 2.17 (0.80-5.87) | 0.128 | - | - |
| Switched ART |  |  |  |  |  |  |
| Yes | 24 (39.3) | 30 (47.6) | 1.4 (0.69-2.86) | 0.353 | 0.89 (0.31-2.53) | 0.827 |
| No | 37 (60.7) | 33 (52.4) | 1 |  | 1 |  |
| HAART initiated |  |  |  |  |  |  |
| TDF based | 27 (44.3) | 12 (19) | 0.3 (0.13-0.66) | 0.003 | 0.26 (0.05-1.27) | 0.096 |
| Non-TDF | 34 (55.7) | 51 (81) | 1 |  | 1 |  |
| HAART on use |  |  |  |  |  |  |
| TDF based | 37 (60.7) | 22 (34.9) | 0.35 (0.17-0.72) | 0.005 | 0.85 (0.25-2.85) | 0.795 |
| Non-TDF | 24 (39.3) | 41 (65.1) | 1 |  | 1 |  |
| Baseline CD4 count (cells/μL) | | | | | | |
| <100 | 20 (32.8) | 17 (27) | 0.76 (0.35-1.64) | 0.481 | 0.86 (0.35-2.11) | 0.745 |
| ≥100 | 41 (67.2) | 46 (73) | 1 | | 1 | |
| WHO clinical stage at interview |  |  |  |  |  |  |
| I/II | 32 (52.5) | 26 (41.3) | 1 |  | 1 |  |
| III/IV | 29 (47.5) | 37 (58.7) | 1.57 (0.77-3.19) | 0.213 | 1.04 (0.44-2.46) | 0.931 |

\* Median age [IQR] ; cOR: Crude odds ratio; aOR: Adjusted odds ratio

## Discussion

The present study aimed at determining acquired HIV drug resistance mutations (HIVDRMs) and predictors of VF in HIV-infected individuals failing to respond to first-line ART in Moshi municipality, northern Tanzania. High profiles of HIVDRMs conferring acquired HIV drug resistance to NRTIs, NNRTIs and PIs were found. Age was a significant factor associated with VF.

In the present study, most (96%) of the successfully sequenced samples had at least one major mutation conferring resistance to HIV drugs. NNRTI resistance mutations profiles (92%) were at the lead compared to NRTIs resistance mutation profiles (68%) consistently with findings elsewhere [6]. The most frequent NNRTI mutations were K103N, V106M, and G190A and they are implicated to confer high level resistance to efavirenz and nevirapine [14]. NNRTIs were previously reported to have low genetic barrier and hence highly vulnerable to resistance [15]. These findings further support the Tanzanian programmatic intervention to replace efavirenz with dolutegravir which has proven to have high genetic barrier to resistance [16]. This study further reports frequent NRTI mutations as M184V, K70R, and D67N in line with other studies [10]. M184V associates with high level resistance to abacavir, emtricitabine, and lamivudine; while K70R and D67N are among thymidine analogue-associated resistance mutations (TAM). More than a half of sequenced samples had at least one TAM. Existence of TAMs reduce the susceptibility of all available NRTIs with an exception of emtricitabine and lamivudine [17].

In addition, this study reports co-existence of T69D mutation with at least one TAM in three samples, two of these samples had M41L and T215Y TAMs, a combination which further fuel cross-resistance to all US FDA approved NRTIs, a phenomenon known as Multi-NRTI resistance [14,18]. This means that the Tanzanian AIDS control program is at risk of remaining with fewer options of main-stream HIV drugs both in first and second-line regimens. Presence of T69 insertions along with TAMs in northern Tanzania further advocates the need for implementation of HIV drug resistance testing and monitoring before and after switching HAART. Continuous programmatic monitoring of HIV drug resistance in HIV-infected individuals failing to respond to first-line HIV drugs will preserve the integrity of few NRTI options in second-line. However there is a need to consider deployment of NRTIs with substantial genetic barrier to resistance.

In the other hand, this study describes full genotypic susceptibility of prescribed HAART to a 49 years old individual with ID K15, on ART for 12 years and with HIV plasma viral load 81,123 copies/ml. At the interview, he reported good adherence to the prescribed HAART. This may be explained by presence of HIV drug resistance mutations away from the *pol* gene and hence necessitating the need for sequencing the whole viral genome in this sample.

A significant association between age and VF concurs with findings from Cameroon [19], Senegal [20], Ethiopia [21], Swaziland [22], Kenya [23,24], Rwanda [25] and Uganda [26]. HIV infected adolescent and young adults on HAART were previously reported to have inconsistence in adhering to antiretroviral medication due to anxiety, depression, forgetfulness, fear of disclosure, ART adverse events and abandoning medication when they feel better [27], the consequences which contributes to VF. Therefore, special care and treatment to the adolescent and young adults is of paramount with emphasis on health education regarding the importance of disclosure and adhering to medications. In addition, assessment and treatment of cognitive and mental health problems is also needed.

Although not statistically significant, this study reports the association between advancement of WHO clinical stage of AIDS and VF, a finding which was consistent with those reported by Jobanputra et al in (2015). Virological failure in individuals with advanced HIV/AIDS can be explained by the severe state of immunodeficiency which attracts opportunistic infections (OIs) and replicative fitness of HIV [28]. Early diagnosis of HIV, effective treatment and monitoring of HIV/AIDS and related OIs to patients on HAART is highly recommended [9].

Tenofovir (TDF) based combination antiretroviral therapy is one of the recommended first-line antiretroviral of choice to the adolescents and adults living with HIV worldwide [9] as well as in Tanzania [8]. Although not statistically significant, being on tenofovir based first-line HAART was protective against VF, this was consistent with studies previously done in the north America [29], Thailand [30] and in the systematic review [31]. On top of that, TDF based regimens were extensively reported in previous randomized clinical trials to be more effective than other combination therapies in bringing favourable virological and immunologic outcomes [32,33]. The possible explanation of this fact is that, compared to other back-borne antiretroviral, TDF based therapy is consumed once daily with minimal treatment related toxicity [32,34], more tolerable [35] and can be easily adhered by PLHIV [36]. Therefore we complement the prescription of TDF based HAART in PLHIV initiating HAART as the strategy of reducing VF.

### Limitations of the study

One of the limitations of this study is to rely on self-reporting of adherence to HAART and pill counting at the clinic visit. It is certain that some of the study participants might have provided imprecise information about adherence. However, this was the common practice prescribed under the Tanzanian HIV and AIDS treatment and management guideline of October 2017. In addition, recall biases are certain to some of the information requested from the VS group, for example it was difficult to report when was the last day to miss taking pills.

Finally, the profiles of HIVDRMs should not be generalized as some of the samples were not sequenced due to financial meltdowns. However, compared to the data in this study, similar high proportions of HIVDRMs are anticipated in the samples yet to be sequenced.

## Conclusion

There is high rate of HIVDRMs in HIV-infected individuals failing to respond to first-line HAART in northern Tanzania. This happens when there is no programmatic monitoring of HIV drug resistance in individuals with VF. Prompt intervention are required to safeguard the limited number of second-line HIV drug options, one of them is implementation of HIV drug resistance monitoring before and after switching HAART. In addition, consideration to deploy HIV drugs with higher genetic barrier is of importance.

## Acknowledgements

The authors are thankful for the permission given by Kilimanjaro Regional Administrative Secretary to conduct research in Mawenzi Regional Referral Hospital, Pasua and Majengo Health Centres. Also many thanks should go to the executive director of Kilimanjaro Christian Medical Centre for allowing us to conduct research in the CTC/CCFCC. The authors are really grateful to the study participants who consented/assented to take part in this study as well as the technical and clinical staffs from Kilimanjaro Christian Medical Centre, Kilimanjaro Clinical Research Institute, Mawenzi Regional Referral Hospital, Pasua and Majengo Health Centre for their support. Extensive gratitude should go to KCRI-Biotechnology research laboratory staff, particularly to Mr. William K. Faustine for their cooperation and technical support during the conduction of this research. Special thanks should go to Professor John Bartlett for critical review of the manuscript. Finally the authors would like to appreciate the HIV Genotyping Facility in Temeke Specialized Laboratory at Temeke Regional Referral Hospital under the Management and Development for Health (MDH, Temeke, Dar es Salaam) for hosting the wet laboratory procedures.

---

S1 Dataset. Demographic and clinical data

